# Optimal Reinforcement Learning with Asymmetric Updating in Volatile Environments: a Simulation Study

**DOI:** 10.1101/2021.02.15.431283

**Authors:** Mojtaba Rostami Kandroodi, Abdol-Hossein Vahabie, Sara Ahmadi, Babak Nadjar Araabi, Majid Nili Ahmadabadi

## Abstract

The ability to predict the future is essential for decision-making and interaction with the environment to avoid punishment and gain reward. Reinforcement learning algorithms provide a normative way for interactive learning, especially in volatile environments. The optimal strategy for the classic reinforcement learning model is to increase the learning rate as volatility increases. Inspired by optimistic bias in humans, an alternative reinforcement learning model has been developed by adding a punishment learning rate to the classic reinforcement learning model. In this study, we aim to 1) compare the performance of these two models in interaction with different environments, and 2) find optimal parameters for the models. Our simulations indicate that having two different learning rates for rewards and punishments increases performance in a volatile environment. Investigation of the optimal parameters shows that in almost all environments, having a higher reward learning rate compared to the punishment learning rate is beneficial for achieving higher performance which in this case is the accumulation of more rewards. Our results suggest that to achieve high performance, we need a shorter memory window for recent rewards and a longer memory window for punishments. This is consistent with optimistic bias in human behavior.

## 1 Introduction

Reinforcement learning (RL) algorithms as normative models present optimal learning rule for the problem of interactive learning in uncertain environments [Sutton and Barto, 2018]. Specially standard RL algorithm or Rescorla-Wagner model [Rescorla, 1972] with a fixed learning rate can follow the changes of reward statistics in a volatile environment. While learning rate in this and other RL algorithms play a role similar to precision-based weighting of prediction error in Bayesian algorithms and so can be updated at each trial. RL algorithms with fixed learning rate are easy to implement and possibly is more common for describing the human and other organisms’ behavior.

In the classical standard RL model, it has been shown that optimal learning rate increases as volatility of environment increases [Behrens et al., 2007, Farashahi et al., 2017]. The change in environment characteristics affects the learning rate of human subjects and it seems that decision maker chooses an appropriate learning rates in different environments with different volatilities. Recent studies show that humans are more likely to use alternative reinforcement learning model instead of classical RL model [Frank et al., 2007, Niv et al., 2015]. In this model subjects weigh the positive and negative prediction error differently and accordingly they have two different learning rates: one for positive prediction errors and another for negative prediction errors.

The dual learning rate model has evidences in human behavior and its underlying neural processing and it is inspired by the bias toward good news versus bad news [Sharot et al., 2007]. This optimism bias helps humans to have better mood and prevent disappointment. But it may have disadvantages in some situations and may hinder optimal decision making. Besides the optimism bias, there is another bias in belief updating toward optimistic news that can be the underlying mechanism for observed bias in humans behavior in RL tasks [Lefebvre et al., 2017, Sharot, 2012]. Moreover, loss aversion is a trait in which decision maker is averse to losing reward relative to gaining an equivalent reward [Tversky and Kahneman, 1991]. Both optimism bias and loss aversion are biases that are prevalent in human decision making. So, the observed different weights for losing and winning in human behavior can be a result of their interplay.

Cognitive flexibility is an essential ability in humans to interact and survive in a complex and volatile environment [Cools, 2019, Cools and Robbins, 2004]. The Probabilistic Reversal learning (PRL) task is a cognitive decision-making task that represents the volatile environment [Pessiglione et al., 2006, Den Ouden et al., 2013, Kanen et al., 2021]. To successfully interact with a complex environment, it is critical to have a trade-off between cognitive stability and cognitive flexibility in order to ignore rare events in stable conditions and flexibly update previous beliefs in the changing conditions. In this study we consider a modified version of a two-option PRL task which is addressed in [Farashahi et al., 2017].

We aim to evaluate whether adding a punishment learning rate to the learning model increase performance in a volatile environment and what is the optimal parameters for reaching the optimal performance. The rest of this paper is organized as follows. Section (2) presents the method which contains behavioral (PRL) task, RL models, and simulation details. Results of Simulation are presented in Section (3). Section (4) discusses the results of simulations and concludes the paper.

## 2 Method

In this study, we aim at simulating the decision-making process during a PRL task. For this purpose we consider a modified version of a two-option PRL task which is addressed in [Farashahi et al., 2017] and simulate it for a wide range of possible environments. We then use RL models to simulate the decision making in each of the designed environments. In what follows we first describe the structure of the PRL task. Then we explain the RL models we use and our simulation procedure.

### 2.1 Probabilistic Reversal Learning task

Reversal learning task represents a volatile environment with uncertainty. In this study we used a modified version of PRL task with two choice options [Farashahi et al., 2017, Den Ouden et al., 2013]. On each trial, two visual stimuli were represented in right or left of the fixation point and participants were instructed to select between two alternative options and receive feedback Fig. 1. Each stimulus was probabilistically associated with reward or punishment. Choosing the mostly rewarded stimulus, called Correct stimulus, delivers reward with the probability of *p*_*Cor*_ and the mostly punished stimulus, Incorrect, results in reward with complementary probability, *p*_*Incor*_ = 1 − *p*_*Cor*_. The level of uncertainty in an environment is determined by *p*_*Cor*_ where the high value indicates a certain environment.

**Figure 1:**
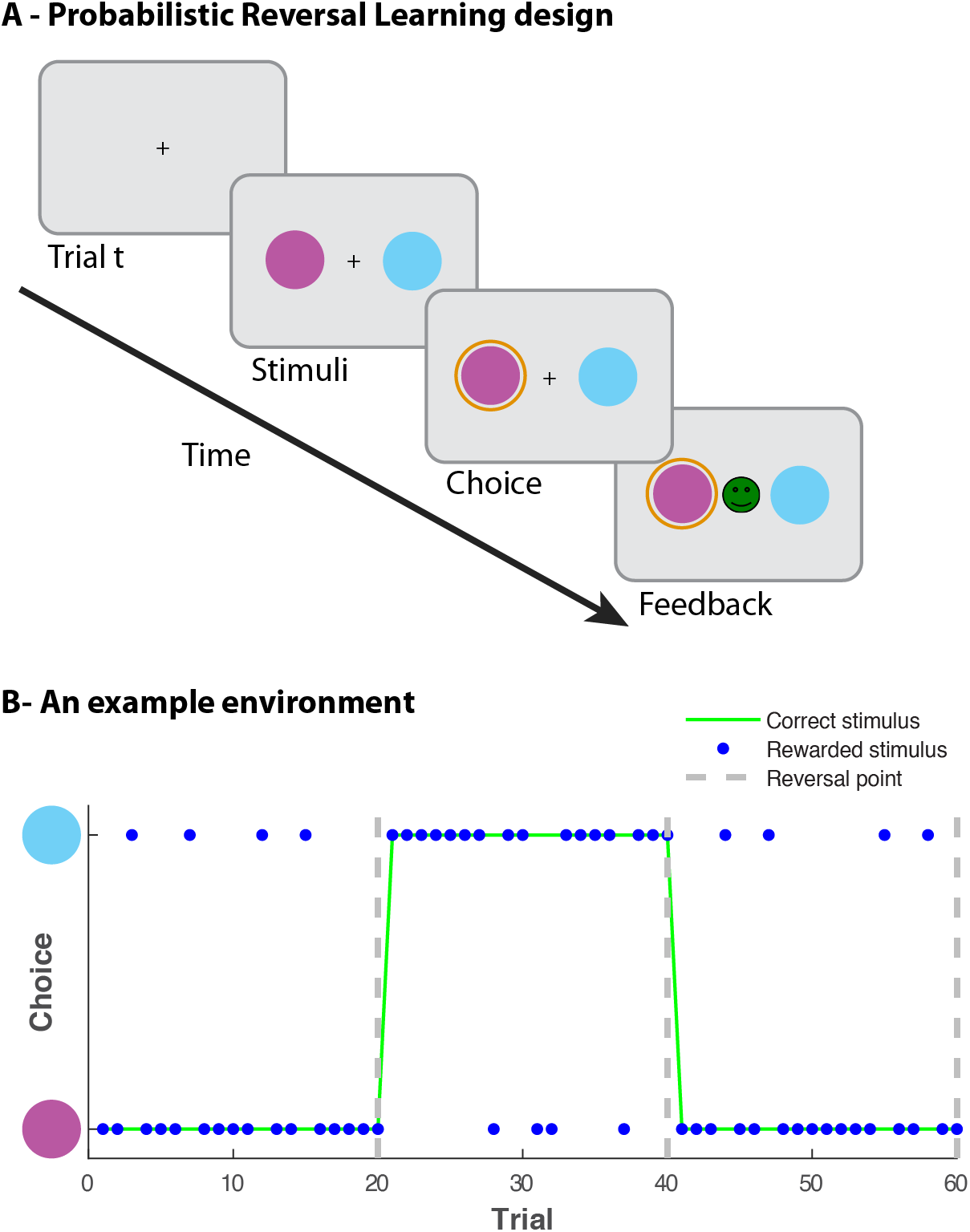
Behavioral task. (A) Probabilistic Reversal Learning task. (B) An example environment with *p*_*Cor*_ = 0.8, *L* = 20 and two reversal points. In each trail, the green line indicates the correct stimulus and blue dots determines the rewarded stimulus. Thus, on the trials that correct and rewarded stimulus are different, subjects received ‘misleading’ feedback due to the environment uncertainty.

After a certain number of trials, *L*, the reward contingencies was reversed, so the mostly rewarded stimulus now became mostly punished and vice versa and this can happen several times. Participants were informed that the contingency rules would change but they were blind to the number of trials between reversals. The level of volatility of environment is defined by *L* and a large one represents a stable environment. Fig. 1B shows an example environment with *p*_*Cor*_ = 0.8 and *L* = 20.

### 2.2 Computational model space

In order to have a good performance in the PRL task, the subject has to continuously update the action values for each of the two options based on receiving feedback. We employed Reinforcement Learning (RL) models as described below.

#### 2.2.1 Rescorla-Wagner (RW) model

The RW model is known as the standard RL model [Sutton and Barto, 2018, Watkins and Dayan, 1992], which contains a single learning rate parameter to update the expected action values, *V*_*c,t*_, based on the environment feedback. The update equation for the expected value is described by the following equation:

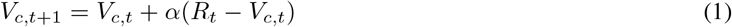

where *α* is the learning rate. The expected value of choice *c* on trial *t*, *V*_*c,t*_, is updated by integrating the environment feedback *R*_*t*_ ∈ {1, −1}. The higher learning rate results in a greater weight on a prediction error, *δ*_*t*_ = *R*_*t*_ − *V*_*c,t*_, thus faster updating in the expected value. On other hand with a lower learning rate, the expected value leads to integration over a wider range of choice-outcome observations. Note that the expected value corresponding to the unchosen option, *V*_~*c,t*_, remains unchanged:

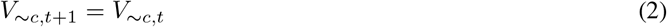

#### 2.2.2 Dual learning rate, Reward-Punishment (RP) model

The RP model is an extended version of the standard RL model [Frank et al., 2007, Den Ouden et al., 2013, Brolsma et al., 2020]. This model utilizes separate mechanisms for learning from positive and negative prediction errors, using dual learning rates. Thus, the expected action value is updated with reward and punishment learning rates for positive and negative prediction error, respectively. The values are updated as follows:

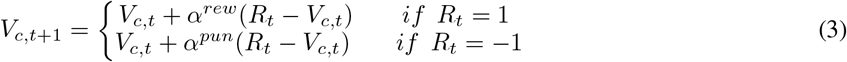

where *α*^*pun*^ is the punishment learning rate and indicates the impact size of negative prediction error on the action value for unexpected punishment. The reward learning rate, *α*^*rew*^, reflects the degree of effect that positive prediction error has on updating the action value. Note that only the value corresponding to the chosen option is updated. In comparison to the RW model, the RP model has one extra parameter, which increases the degree of freedom and can results in more flexible behavior.

For both models, to select an action based on the computed values, a soft-max choice function was employed to calculate the probability of each choice, *left* or *right*. For the current PRL task, *j*∈ {*left, right*} is the set of all possible options and the probability is computed as follows:

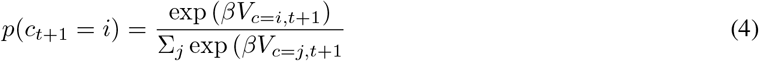

where *β* is the inverse temperature parameter, also known as the exploration/ exploitation parameter, which reflects the stochasticity of choices. A high *β* means choosing the option with higher expected value more consistently.

#### 2.2.3 Simulation

In order to quantify and evaluate the performance of each model, we performed an extensive simulation, which is the first and important step of each computational neuroscience study [Wilson and Collins, 2019]. As described above, the PRL task environment is determined by two parameters, the level of uncertainty *p*_*Cor*_ and the level of volatility *L*. First, we created different environments using all possible combinations of the following parameter values *p*_*Cor*_ = [0.6, 0.8] with a step size of 0.02, and *L* = [20, 200] with step size of 20. As we aimed to run 2000 trials for each environment, the number of reversals are different for environments with different level of volatility. The environment feedback for each stimulus type is generated as a pseudo-random series based on *p*_*Cor*_.

Then, for each of 110 environments, we computed the maximum performance that can be reached using both models. The free parameters for RW model are *θ*_*RW*_ = {*α, β*}, and for RP model are *θ*_*RP*_ = {*α*^*rew*^, *α*^*pun*^, *β*}. In order to find the optimal values for *α*, *α*^*rew*^, and *α*^*pun*^ we simulated performance score for the entire [0, 1] range of these three parameters with *step size* = 0.05. For *β* parameter we used the median value reported in other studies, *β* = 4.5 [Den Ouden et al., 2013, Kandroodi et al., 2020]. Although we checked the consistency of the results for *β* = [2, 8]. The summary of the parameters detail for simulation is reported in Table 1.

**Table 1:** Summary of the parameters detail for simulation. It contains environmental and models parameter

| Parameter | Range | Step size | #Sim. values | Type of parameter |
| --- | --- | --- | --- | --- |
| $L$ | [20, 200] | 20 | 10 | Environmental |
| $p_{Cor}$ | [0.6, 0.8] | 0.2 | 11 | Environmental |
| $\alpha$ | [0, 1] | 0.05 | 21 | RW model |
| $\alpha_{rew}$ | [0, 1] | 0.05 | 21 | RP model |
| $\alpha_{pun}$ | [0, 1] | 0.05 | 21 | RP model |

We simulated 20 artificial agents for each parameter combination set and reported the average (normalized) performance of these agents. We computed a normalized performance score, *S*_*p*_, based on the likelihood of receiving positive feedback for each of the stimulus options [Kandroodi et al., 2020]:

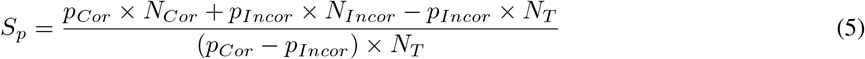

where *N*_*Cor*_ (*N*_*Incor*_) is the number of trials in which Correct (Incorrect) stimulus is selected and *N*_*T*_ is the total number of trials. The last term in the numerator and the term in the denominator is added to normalize the performance score to a minimum of 0 when the incorrect stimulus was chosen on all trials, and a maximum of 1 when the correct stimulus was chosen on all trials. The normalized performance score is comparable across different environment.

## 3 Results

We generated the simulation data for 110 environments with different levels of uncertainty and volatility. The behavior in each environment was therefore simulated for 21^2^ = 441 combination of free parameters {*α*^*rew*^, *α*^*pun*^} for RP model and 21 value of learning rate for RW model, see Table 1. In order to reduce noise in behavior we simulated 20 artificial agents for each parameter combination set and reported the average performance of these agents. In what follows we first report the optimal performance of the models in different environments and their comparison. Then we present the optimal learning rate for the RP model.

### 3.1 Optimal performance

The landscape for optimal performance is demonstrated in Fig. 2. The key observation is that the optimal performance is lower for the more volatile (small *L*) and more uncertain (small *p*_*Cor*_) environments. In contrast, it was higher for the more stable (big *L*) and more certain (big *p*_*Cor*_) environments. This trend is consistent for both models, Fig. 2A-B.

**Figure 2:**
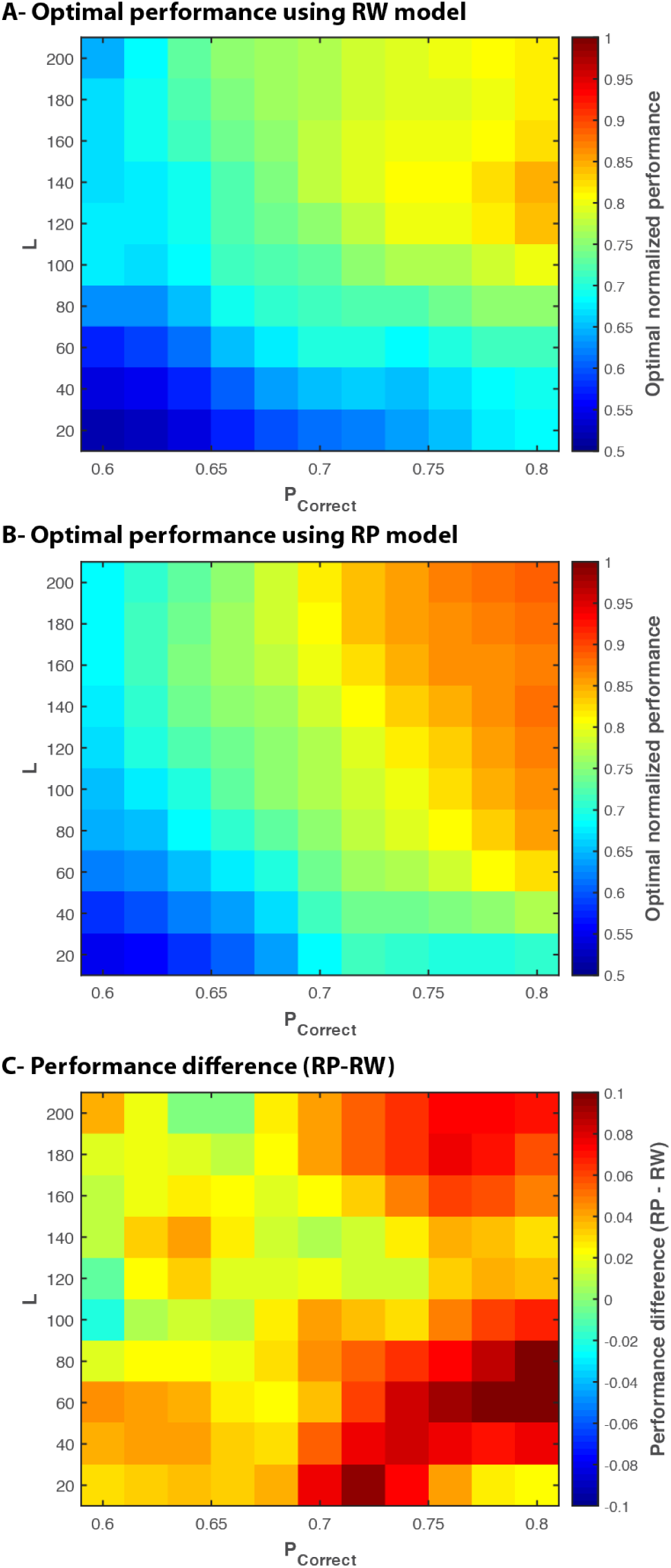
The optimal performance score landscape as a function of environment variables: level of uncertainty (*p*_*Cor*_) and level of volatility (*L*). The normalized performance of 0.5 indicates the chance level. Optimal performance corresponding to the environment and the model is computed by averaging top 2% performance in order to reduce noise (see [Farashahi et al., 2017]). (A) The optimal performance using RW model. The optimal performance is lower for the more volatile (small *L*) and more uncertain (small *p*_*Cor*_) environments. In contrast, it is higher for the more stable (big *L*) and more certain (big *p*_*Cor*_) environments. (B) The optimal performance using RP model. (C) The difference between optimal performance of RP and RW model. The RP model shows better optimal performance in different environments. On average, the RP model has 0.04 higher performance than RW model.

The difference between optimal performance of the two models (RP-RW) is presented in Fig. 2C. The RP model performed better, average difference=0.04, than RW in almost all environments and this difference is bigger for more certain environments.

### 3.2 Optimal learning rate

The optimal learning rate for the RP model is computed based on the set of parameters that resulted in the best performance in each of the environments. The landscape for optimal reward learning rate is demonstrated in Fig. 3A. The figure shows that optimal reward learning rate is higher both for the more volatile but more certain (bottom right corner) and the more uncertain but stable (top left corner). It can also be seen that when environment is more stable and certain (top right corner), the optimal performance is reached by comparably lower reward learning rate. The landscape for optimal punishment learning rate is demonstrated in Fig. 3B. As a general trend, a smaller punishment learning rate is needed for optimal behavior in the more certain environments. The reward learning rate to punishment learning rate ratio is demonstrated in Fig. 3C. Notice that, for reward learning rate equal to punishment learning rate, *α*^*rew*^/*α*^*pun*^ = 1, the RP model reduces to the RW model. The results indicate that in almost all environments, reaching the optimal performance needs the reward learning rate to be greater than the punishment learning rate. The black line shows the level contour equal to one, where the optimal performance occurs at the reward learning rate equal to the punishment learning rate. For the rest, optimal performance occurs when the reward learning rate is higher than the punishment learning rate.

**Figure 3:**
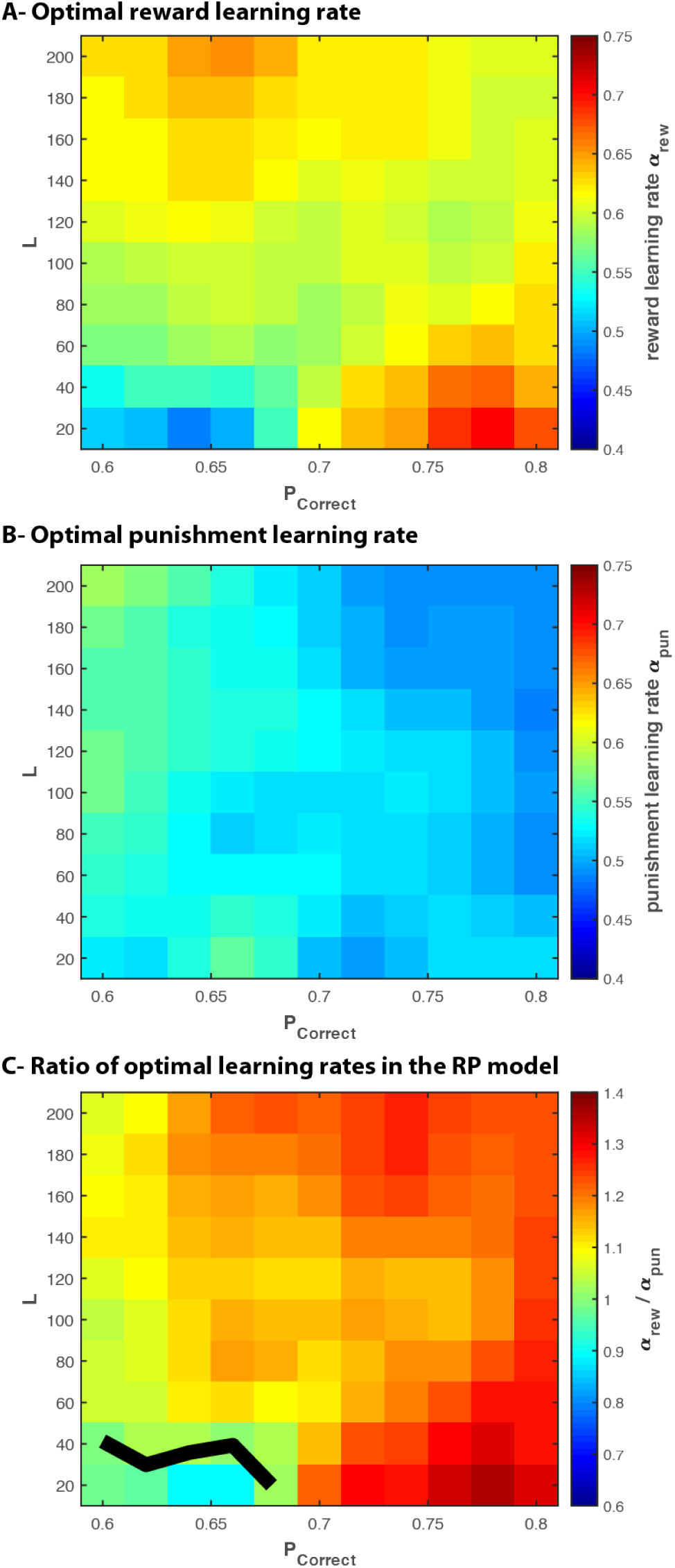
The optimal learning rates landscape for the RP model as a function of environment variables: level of uncertainty (*p*_*Cor*_) and level of volatility (*L*). (A) The optimal reward learning rate. The optimal reward learning rate is higher both for the more volatile but more certain (bottom right corner) and the more uncertain but stable (top left corner). The optimal performance is reached by comparably lower reward learning rate when the environment is more stable and certain (top right corner). (B) The optimal punishment learning rate. A smaller punishment learning rate is needed for optimal behavior in the more certain environments. (C) The ratio of the optimal learning rates for the RP model. The results demonstrates that for almost all environments, reaching the optimal performance needs the reward learning rate to be greater than the punishment learning rate. The black line shows the level contour, where the optimal performance occurs at the reward learning rate equal to the punishment learning rate.

## 4 Discussion and conclusion

In this study, we aimed at finding the optimal learning rates in a reinforcement learning model with two different learning rates for positive and negative prediction errors. The optimal performance for different learning rates show that using two different learning rates results in a better performance compared to single learning rate. The results demonstrate that for almost all environments, reaching the optimal performance needs the reward learning rate to be greater than the punishment learning rate.

Our simulation results are aligned with the study of [Lefebvre et al., 2017] in which it is shown that human decision makers have higher reward learning rate than punishment learning rate. Our results can be a base for justification of optimistic update bias [Sharot et al., 2011] and introduce an advantage for this bias in volatile environments. From evolutionary perspective, it seems that as organisms encountered high volatile environments, they possibly developed optimism in belief updating.

It is important to note that higher learning rate does not mean higher learning, instead it shows the backward extension of memorizing rewards. In other words, learning rate somehow shows the relative weight of recent versus old rewards; i.e. higher learning rates put emphasis on the recent reward while lower learning rates use older rewards relatively more. It is a kind of forgetting factor, and as learning rate increases older reward are forgotten more. So, higher reward learning rates show shorter integration window for rewards, while smaller punishment learning rates show longer integration window. Thus, optimal strategy in our simulation emphasizes recent rewards more and integrates punishments in longer windows. Optimal strategy has longer memory of punishments and failures and shorter memory for recent rewards.

There are studies that have already investigated the fitting of dual learning rate RL model to the behavior and higher reward learning rate seems to be valid [Sharot et al., 2007, Lefebvre et al., 2017]. However, it is important to remind that the application of this model for behavioral studies in volatile environments and observing the learning rates of human subjects need to be investigated in future studies. Moreover, instead of constant learning rates, using some adaptive strategies to adjust learning rates according to environment statistics is an open question. Future studies need to find a way to estimate environment volatility and use them in adjustment of learning rates. Some models in Bayesian modeling, such as Hierarchical Gaussian Filtering (HGF) [Mathys et al., 2014] provide Bayesian insights on this issue.

Optimal performances in different environments show that reward accumulation expectedly increases as environment become more stable and more certain. Importantly, added benefit of having two learning rates is small in environments that are uncertain, and performance improvement increases for environments with certainty irrespective of volatility. Optimal punishment learning decreases as the environment becomes more certain, and so higher punishment learning rates are needed for uncertain environments. Optimal reward learning rate shows an interaction between volatility and uncertainty. The highest reward learning rates are obtained both for the more certain but volatile and the more stable but more uncertain environments. These all show that optimal learning strategy can be adjusted depending on the statistics of environment.

In conclusion, by knowing the statistics of the environment, both uncertainty and volatility, we can use obtained parameters for the accumulation of more reward. The proposed strategy seems to have a higher reward learning rate and so short memory window for recent rewards and a longer memory window for punishment learning rates.

